# Graphene-enabled, spatially controlled electroporation of adherent cells for live-cell super-resolution microscopy

**DOI:** 10.1101/642868

**Authors:** Seonah Moon, Wan Li, Ke Xu

## Abstract

The incorporation of exogenous molecules into live cells is essential for both biological research and therapeutic applications. In particular, for the emerging field of super-resolution microscopy of live mammalian cells, reliable fluorescent labeling of intracellular targets remains a challenge. Here, utilizing the unique mechanical, electrical, and optical properties of graphene, a single layer of bonded carbon atoms, we report a facile approach that enables both high-throughput delivery of fluorescent probes into adherent live cells and *in situ* super-resolution microscopy on the same device. ∼90% delivery efficiencies are achieved for free dyes and dye-tagged affinity probes, short peptides, and whole antibodies, thus enabling high-quality super-resolution microscopy. Moreover, we demonstrate excellent spatiotemporal controls, which, in combination with the ready patternablity of graphene, allow for the spatially selective delivery of two different probes for cells at different locations on the same substrate. We thus open up a new pathway to the microscopic manipulation and visualization of live cells.

---

The incorporation of exogenous molecules into live cells is essential for both biological research and therapeutic applications^1–3^. For example, in bioimaging, it is often necessary to deliver exogenous genes or probes into cells to visualize the targets of interest. In particular, for the emerging field of super-resolution microscopy (SRM) of live mammalian cells^4–8^, although the intracellular gene expression of fluorescent proteins (FPs) provides a relatively straightforward approach^4^, FPs offer limited brightness and photostability when compared to organic dyes^9^. On the other hand, high-performance dye-based probes for SRM often do not readily cross the strong barrier created by the cell plasma membrane^6,8^, and so rely on membrane-disruption techniques for intracellular delivery.

Although chemical permeabilization, including the use of mild detergents and toxins^2,10-12^, provides relatively easy means of intracellular delivery, recovery of membrane integrity is often challenging. Microinjection provides a controlled means for intracellular delivery^2^, but is limited in throughput and highly dependent on the operator’s skill. Electroporation creates small, resealable pores on the cell membrane via applied electric fields, and is nowadays routinely used for intracellular delivery owing to its high efficiency and low cell toxicity^2,3,13^. However, electroporation is typically performed for detached and (re-)suspended cells. For the delivery of external fluorescent probes, the long (∼10 h) subsequent re-plating time^6,8^, which is essential for the cells to re-adhere to the coverslip for high-resolution imaging, is inconvenient and potentially gives rise to undesired side effects due to the prolonged introduction of probes. Recent advances in nanotechnology and microfluidics have led to the exciting development of numerous new intracellular-delivery methods,^1,2,14–20^ each overcoming certain limits of traditional approaches. However, so far these technologies have not been designed to enable high-resolution microscopy on the same device after high-throughput delivery.

Here we introduce a graphene-based, facile approach for the direct, high-throughput delivery of fluorescent probes into adherent cells to enable *in situ* live-cell SRM on the same device within minutes. Recent years have witnessed rising research interest in interfacing graphene with cell biology^21–25^. We previously demonstrated the use of graphene to encapsulate wet mammalian cells to enable facile electron microscopy^26^. Here, we utilize the unique properties of graphene to enable cell culturing, electroporation-based probe delivery, and *in situ* SRM imaging on the same device. Besides achieving high delivery efficiencies and consequent high-quality SRM, we further demonstrate excellent spatial and temporal controls, thus realizing the spatially selective delivery of different probes into cells at different parts of the same substrate.

Monolayer graphene, as produced by chemical vapor deposition on copper foils^27^, was deposited onto regular glass coverslips as ∼10×5 mm^2^ pieces, sealed with a small plastic tube, and contacted at both edges (Fig. 1a and Supplementary Fig. 1). Two-point measurements showed resistances of a few kΩ across the as-prepared devices. Interference reflection microscopy (IRM)^28^ was employed to confirm that graphene in the final devices to be continuous monolayers with minimal defects. Adherent mammalian cells (A549 and PtK2 cells) were cultured on the graphene surface under standard tissue culture conditions. Previous work has shown the graphene surface to be suitable for cell growth^24,25,29,30^. We similarly found cells grew well on our graphene electrodes, and further found that the measured graphene resistance increased moderately to ∼10 kΩ after ∼1 day growth (Fig. 1c).

**Figure 1.**
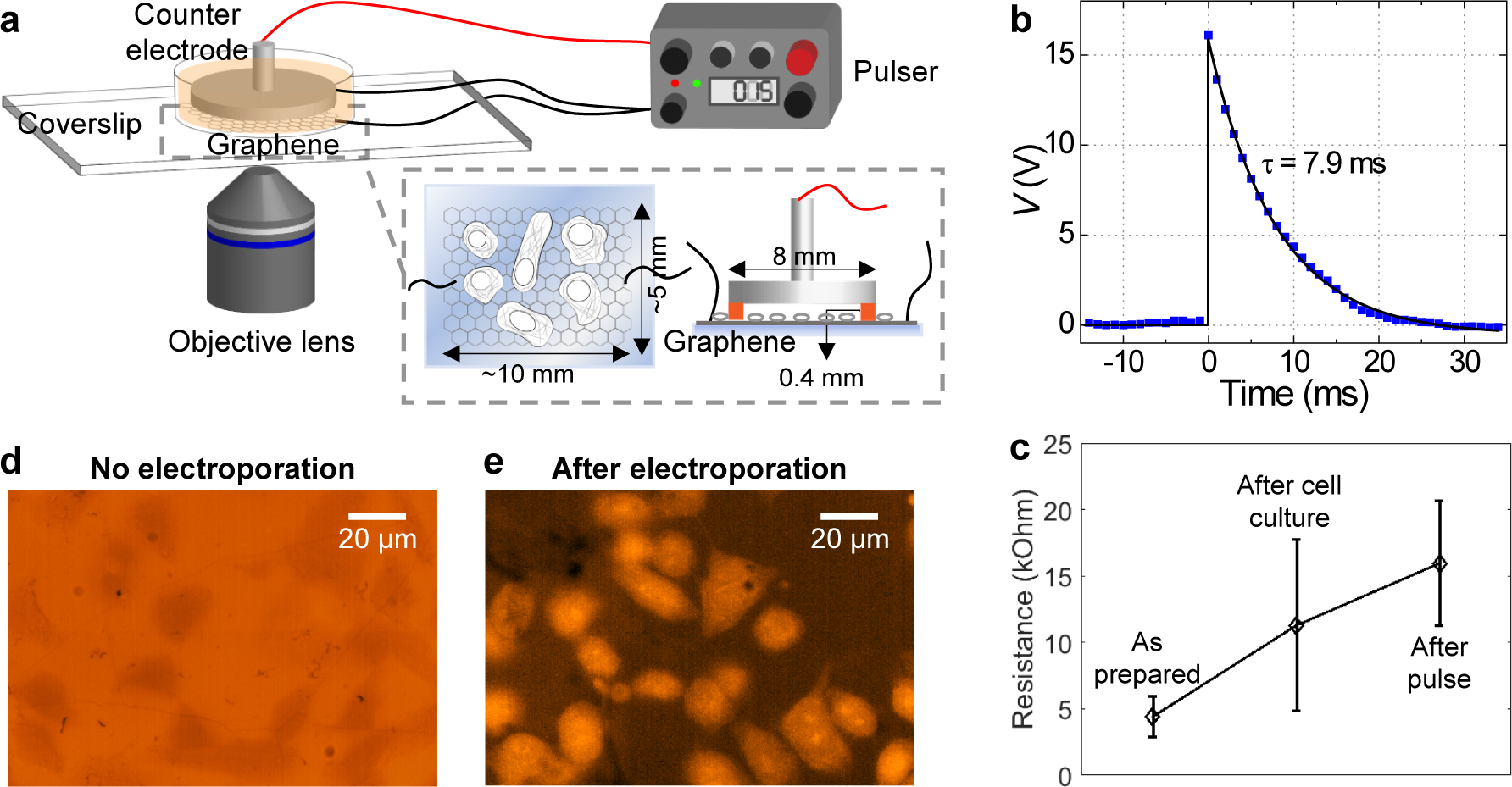
Electroporation of adherent cells on a graphene-covered glass coverslip. (a) Schematic of the experimental setup. (b) A representative pulse shape, as measured from an oscilloscope. (c) Measured two-point resistance of graphene for the as-prepared device, after culturing of adherent cells, and after electroporation (*N* = 4 measured devices). Error bars: standard deviation between samples. (d) Fluorescence micrograph of PtK2 cells incubated in an SR101-containing medium for 10 minutes; cells appear as darker regions due to the physical exclusion of the dye. (e) Fluorescence micrograph of PtK2 cells on graphene after the application of the voltage pulse, 10 min incubation, and wash-out of free dyes.

At the time of probe delivery, the culture medium was replaced with a commercial electroporation buffer solution, to which we added the desired fluorescent probes. Counter electrode was made out of a metal pin-stub mount commonly used in scanning electron microscopy (SEM), which was mounted up-side-down with the application of ∼400 µm-thick Teflon tape on the rim as an insulating spacer (Fig. 1a and Supplementary Fig. 1). This counter electrode was gently placed into the device, and the assembled device was mounted onto an inverted fluorescence microscope. Scanning the microscope focus indicated that the distance between the graphene and counter-electrode surfaces was ∼500 µm. For electroporation, we applied a voltage pulse across the graphene and counter electrodes through capacitor discharging using a commercial electroporator. Efficient probe delivery was readily achieved at low voltages (∼15 V) using a small (10 µF) capacitor, with typical pulse halftimes of ∼5-10 ms (Fig. 1b). A mild increase in graphene resistance was noted after the electroporation process (Fig. 1c), suggesting that the graphene electrode remained mostly intact.

We started with the delivery of a free organic dye, sulforhodamine 101 (SR101, molecular weight: 606.7). As SR101 is not normally taken up by PtK2 cells, microscopy of cells in a medium containing this probe showed lower fluorescence for cell-occupied areas owing to physical exclusion (Fig. 1d). After electroporation in the graphene device, 10 min of incubation, and then washout of free dyes, we found the cytoplasm of most cells became fluorescent due to the incorporation of SR101 (Fig. 1e).

We next examined dye-tagged probes that bind to specific intracellular targets, starting with phalloidin-Alexa Fluor 488 (phalloidin-AF488), a small (∼1.3 kDa), highly specific fluorescent marker for the actin cytoskeleton. As expected, the probe itself was non-permeable for A549 cells, and so cells initially appeared as darker regions in fluorescence micrographs due to physical exclusion (Fig. 2a). After the application of an electroporation voltage pulse, the cells quickly lightened up in ∼1 min (Fig. 2bd), and continued to rise in fluorescent signal over time (Fig. 2cd). Low-magnification images showed, over large areas, most cells on the graphene surface to be efficiently labeled, whereas in the same view, cells on the bare glass substrate remained unlabeled (Fig. 2i).

**Figure 2.**
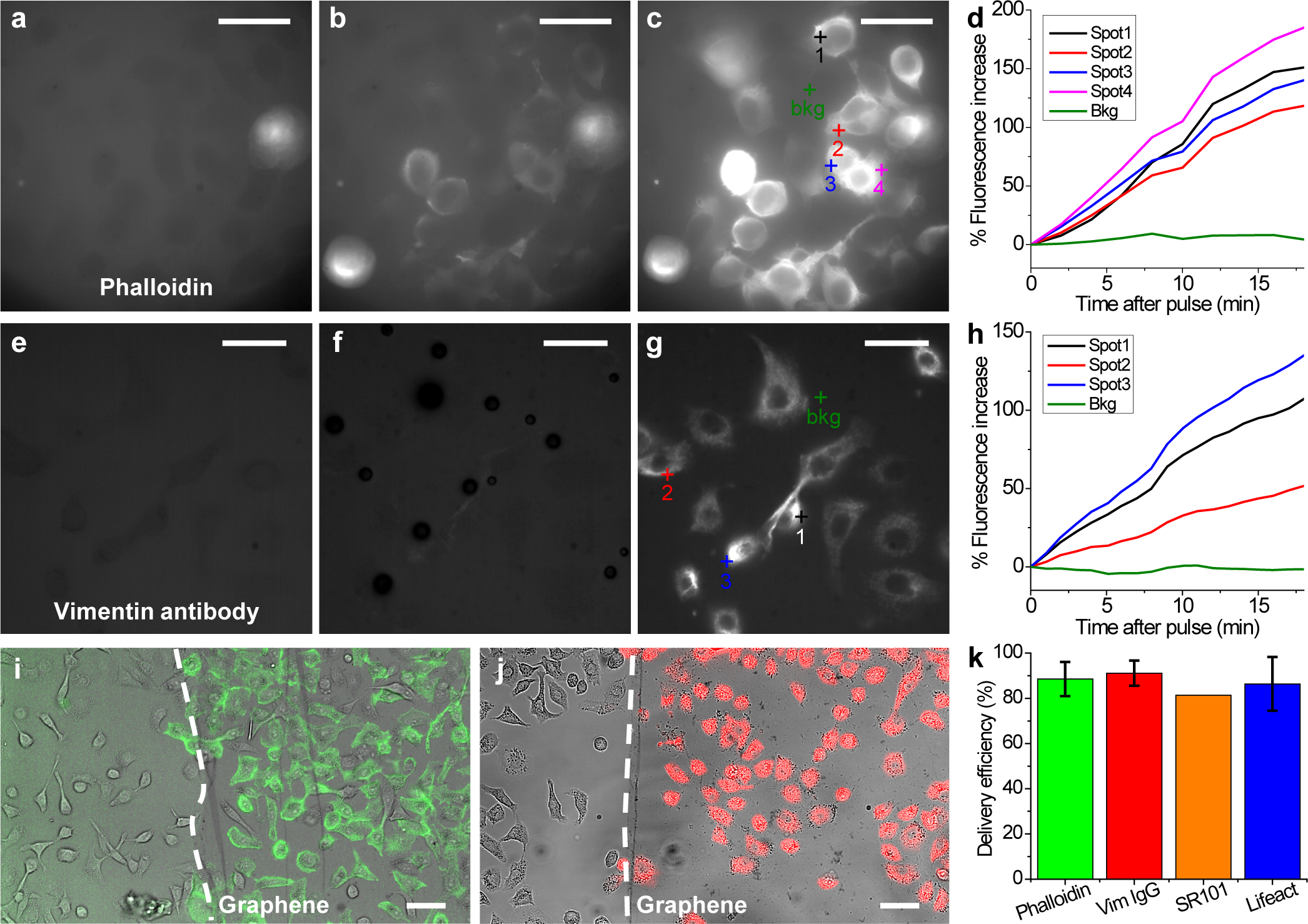
Delivery of dye-tagged probes that bind to specific intracellular targets. (a-c) Delivery of phalloidin-AF488 into A549 cells through graphene electroporation. (a) Fluorescence micrograph of the sample before the application of the voltage pulse. (b,c) Fluorescence micrographs for the same area 1 min (b) and 20 min (c) after the voltage pulse. (d) Change in local fluorescence intensity over time after the voltage pulse, for the different spots marked in (c). (e-g) Delivery of AF647-conjugated anti-vimentin full IgG antibody into A549 cells. (e) Fluorescence micrograph before the voltage pulse. (f,g) Fluorescence micrographs for the same area 1 min (f) and 20 min (g) after the voltage pulse. (h) Change in local fluorescence intensity over time after the voltage pulse, for the different spots marked in (g). (i) Merged transmission (grayscale) and fluorescence (green) micrographs for the spatially controlled delivery of phalloidin in A549 cells adhered to graphene (to the right of the dashed line) vs. no delivery to cells on the bare glass surface without graphene (left of the dashed line). (j) Similar results for the spatially controlled delivery of anti-vimentin IgG (red). (k) Percentage of cells being labeled for phalloidin (*N* = 7 devices), anti-vimentin IgG (*N* = 4 devices), SR101 (*N* = 1 device), and Lifeact (*N* = 4 devices), as determined from correlated transmission and fluorescence micrographs. Error bars: standard deviation between samples. Scale bars: 20 µm (a-c;e-g); 50 µm (i,j).

We next turn to the more challenging task of dye-tagged whole immunoglobulin G (IgG) antibodies. Whereas antibody-based fluorescence labeling (immunofluorescence) is routine for SRM of fixed and permeabilized cells and is favorable for its ease and versatility^8,9^, its use in live-cell microscopy and SRM has been rare^12^ due to difficulties in delivering the sizeable (∼155 kDa) IgG molecules into the cell. We found graphene-based electroporation enabled efficient delivery of dye-tagged IgGs, *e.g.*, Alexa Fluor 647 (AF647)-tagged IgG against vimentin (Fig. 2e-g), although the increase in intracellular fluorescence was slower (Fig. 2e-h) when compared to that of phalloidin, possibly attributable to slower diffusion of the heavy IgGs. Low-magnification images showed that similar to that with phalloidin, highly efficient and selective labeling was achieved for cells on the graphene surface (Fig. 2j). Correlating transmission and fluorescence micrographs showed that, consistently, ∼80-90% of cells on the graphene surface to be successfully labeled (Fig. 2k) for the cases of phalloidin, anti-vimentin IgG, SR101, as well as Cy5-tagged Lifeact, a 17-amino-acid peptide that reversely binds to actin filaments^31^ (see also Fig. 3hi below). Together, we have shown that our graphene-based electroporation allowed for the intracellular delivery of small to large non-permeable probes into adherent cells with high efficiency and good spatiotemporal control.

**Figure 3.**
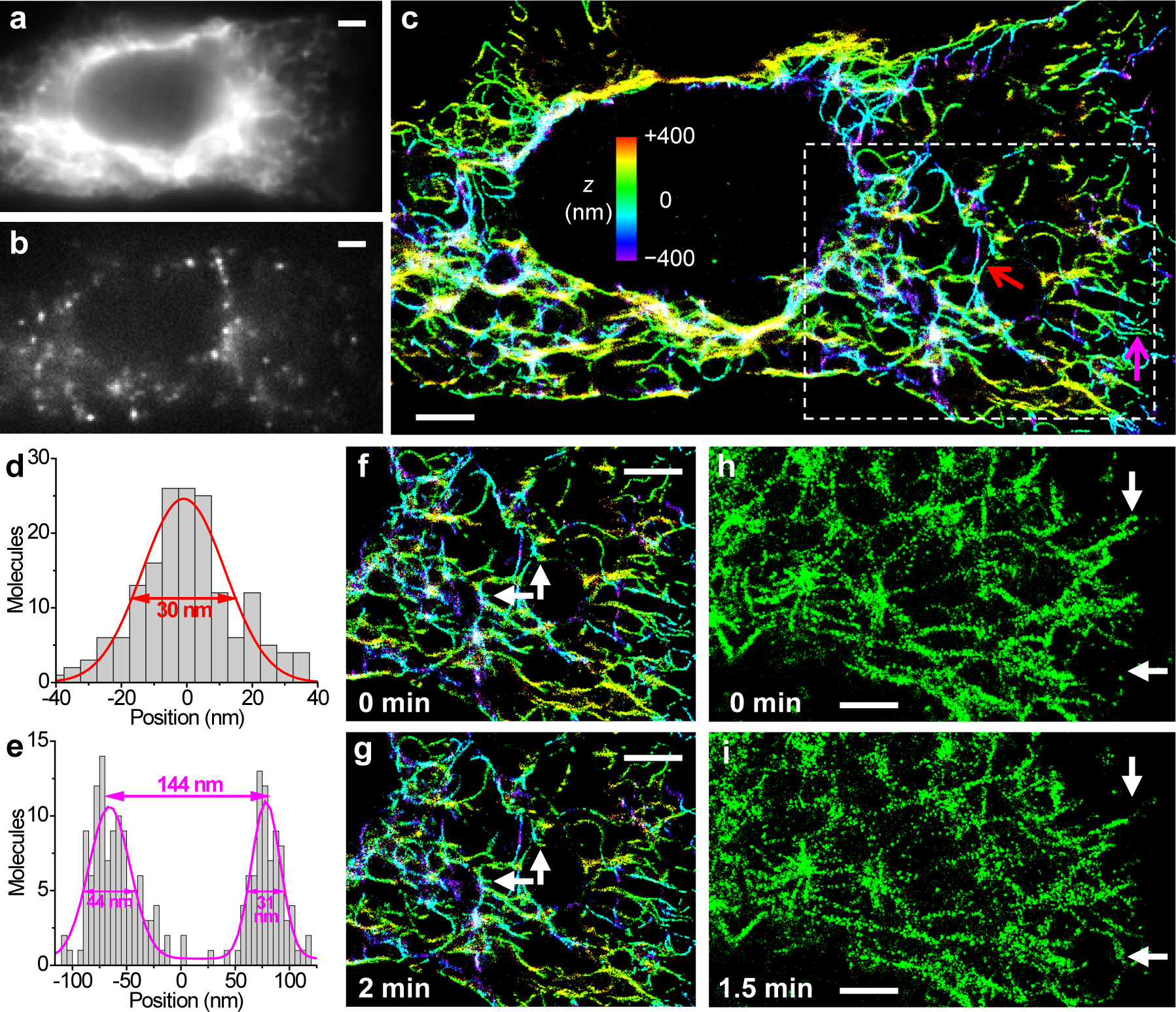
STORM SRM through fluorescent probes delivered by graphene electroporation. (a) Diffraction-limited image of vimentin in a live A549 cell labeled through the graphene-electroporation delivery of an AF647-tagged IgG antibody. (b) A typical frame of single-molecule images during STORM acquisition. (c) Resultant 3D-STORM image. Color presents the depth (z) information (color scale bar). (d) Cross-sectional profile of the single vimentin filament pointed to by the red arrow in (c). Fit to a normal distribution gave FWHM of 30 nm. (e) Cross-sectional profile for two adjacent filaments pointed to by the magenta arrow in (c). (f,g) A sequence of two STORM images at 0 min (f) and 2 min (g). Arrows point to structural changes at the nanoscale. (h,i) A sequence of two STORM images of actin filaments in a live A549 cell labeled through the graphene-electroporation delivery of Lifeact-Cy5, at 0 min (h) and 1.5 min (i). Arrows point to notable structural changes. Scale bars: 2 µm.

Based on the above good delivery results, we next demonstrate the use of our system for *in situ* live-cell STORM (stochastic optical reconstruction microscopy)^32,33^ SRM in the same device immediately after probe delivery. Here we utilize the unique properties of graphene that although it is highly conductive (to enable electroporation), it is only a single layer of atoms and is highly transparent to light, so that we could perform high-resolution microscopy directly through the graphene electrode with minimal issues. Thus, after electroporation delivery of STORM-compatible probes, we replaced the cell medium with a live-cell STORM imaging buffer^6^, and mount the device on a 3D-STORM system equipped with a 100x, oil-immersion objective lens.

Conventional epi-fluorescence images taken at low illumination powers, *e.g.*, for vimentin filaments labeled in live A549 cells through the graphene-electroporation delivery of AF647-tagged IgG, showed no signs of distortion as we imaged through the graphene electrode (Fig. 3a). By next increasing the illumination power, we photoswitched most of the labeled probe molecules into a non-emitting dark state. The reversible photoswitching of these molecules between the dark and emitting states led to well-resolved, bright single-molecule fluorescence (Fig. 3b) that “blinked” stochastically in space and time, which we recorded continuously at 110 frames per second using an EM-CCD. An average of ∼3,500 photons were collected for each emitting single molecule, in agreement with that typically obtained with AF647 in live-cell STORM^6^. Accumulating the 3D localizations^33^ obtained from 47,128 frames of single-molecule images led to 3D-STORM SRM images of high resolution (Fig. 3c). For the single vimentin intermediate filament pointed to by the red arrow in Fig. 3c, cross-sectional profile gave an FWHM (full width at half maximum) width of 30 nm (Fig. 3d), consistent with a convolution of the ∼20 nm spatial resolution of STORM^6,32,33^ with a ∼20 nm diameter of IgG-labeled vimentin filament. Figure 3e further shows a case (magenta arrow in Fig. 3c) in which two filaments are clearly resolved at a center-to-center distance of 144 nm, well below the diffraction limit. Subdividing the collected frames of single-molecule images to construct a sequence of STORM images further enabled the scrutiny of nanoscale structural changes over time (Fig. 3fg). Good live-cell STORM SRM was also achieved for the actin cytoskeleton labeled by Lifeact-Cy5 (Fig. 3hi) and phalloidin-AF647 (Supplementary Fig. 2).

We next further exploit the spatial and temporal control of our graphene-based approach to uniquely enable patterned delivery of two different probes. As a first demonstration, we made a scratch at the center of graphene to divide it into two (top and bottom) halves, which were each separately contacted by a metal wire (Fig. 4a). IRM^28^ (Fig. 4b-d) showed that the scratch was ∼20 µm-wide, for which region graphene was fully removed, and conductance measurements indicated that the two halves were electrically isolated. After plating cells, we first replaced the culture medium with an electroporation buffer that contained AF647-conjugated anti-vimentin IgG, and applied a 15 V pulse only to the bottom half of the graphene electrode against the counter electrode. After ∼15 min incubation, the cells were allowed to recover in a medium containing 2 mM ATP and 2% glucose at 37 °C for ∼20 min^12^. The medium was replaced by another electroporation buffer that contained phalloidin-CF568, and then another 15 V pulse was applied across the top half of the graphene electrode and the counter electrode. Remarkably, the above sequential electroporation procedure enabled patterned delivery, so that fluorescent micrographs taken in the AF647 and CF568 channels showed that the former was selectively delivered into cells on the bottom half of the graphene electrode (Fig. 4e-g;k-m), whereas the latter was selectively delivered into cells in the top half of the device (Fig. 4h-m). Interestingly, we further found that the spatial specificity for the second probe relied on the proper recovery (sealing) of the plasma membrane after the first electroporation step. Skipping this recovery step led to nonspecific delivery into the unsealed cells due to the first electroporation pulse (Supplementary Fig. 3).

**Figure 4.**
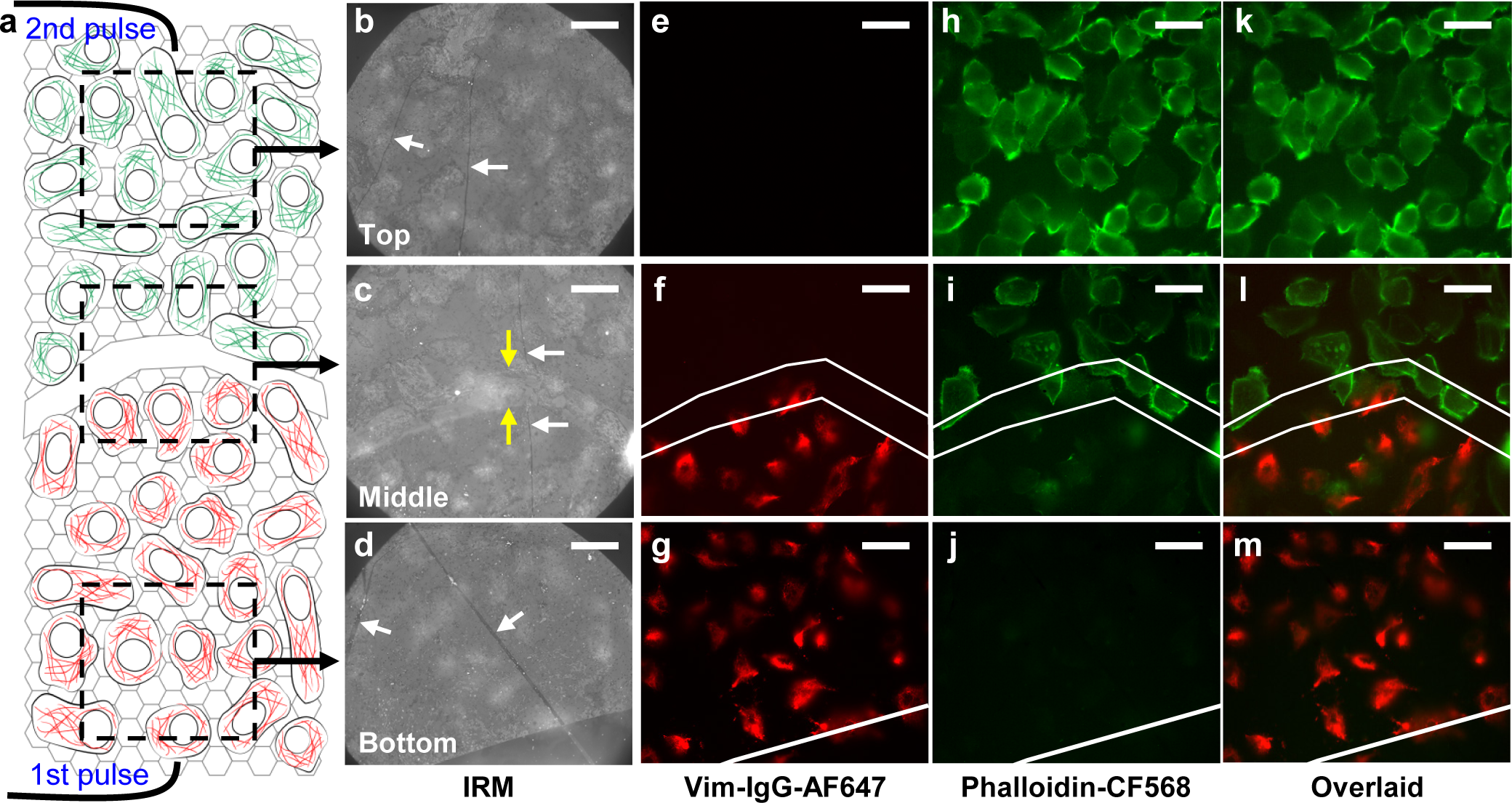
Patterned delivery of two different probes for cells adhered to different regions of the same substrate. (a) Schematic of the sample, with a scratch through the graphene electrode that divided it into two halves. A voltage pulse was first applied to the bottom half in the presence of the first (red) fluorescent probe, cells were recovered for ∼20 min, and then a second voltage pulse was applied to the top half in the presence of the second (green) fluorescent probe. (b-d) IRM images of A549 cells on graphene for the top (b), middle (c), and bottom (d) parts of the sample, as schematized in (a). White arrows point to wrinkles in graphene, which give high IRM contrast. Yellow arrows point to edges of the scratch. (e-g) Fluorescence micrographs for the same areas as (b-d), for the channel of the first fluorescent probe (AF647-tagged anti-vimentin IgG antibody). (h-j) Fluorescence micrographs for the same areas for the second fluorescent probe (phalloidin-CF568). (k-m) Overlay of the two color channels. White lines in (f,i,l) mark the edges of the top and bottom halves of graphene. Scale bars: 20 μm.

In summary, we have developed an integrated system that enables the facile electroporation delivery of fluorescent probes into adherent mammalian cells for immediate single-molecule detection and SRM on the same platform. High (∼90%) delivery efficiency was achieved with low pulse voltages for from free dye molecules up to full IgG antibodies, and the outstanding optical properties of graphene enabled SRM using an oil-immersion objective. Moreover, we demonstrated unique spatial and temporal controls, achieving patterned delivery of two probes for different regions of the same substrate with high selectivity. By removing the need to detach and then re-adhere the cells to coverslips for high-resolution microscopy, as required by typical electroporation methods, our *in situ* approach greatly expedites labeling and reduces the potential adverse effects due to prolonged retention of external probes inside live cells. Furthermore, whereas in this work we have focused on probes based on organic dyes for their ease of visualization, our approach may also enable the delivery of other probes or chemicals, including drugs, into live cells. By being able to deliver different chemicals to different, spatially predefined subsets of cells on the same substrate under the same conditions, a well-controlled, multiplexed platform may thus be constructed for the quantitative examination of drug effects through high-resolution microscopy.

## Acknowledgments

We thank Meghan Hauser and Limin Xiang for assistance and discussion. K.X. is a Chan Zuckerberg Biohub investigator, and acknowledges support from the Bakar Fellows Award and the Packard Fellowships for Science and Engineering. S.M. acknowledges support from Samsung Scholarship.

